# MRTX1133 is a potent non-covalent KRAS (G12C) inhibitor with tissue-specific activity

**DOI:** 10.1101/2025.09.09.674973

**Authors:** Abraham C. Sianoya, Yan Zuo, Cynthia V. Pagba, Jeffrey A. Frost, Paul Mark B. Medina, Alemayehu A. Gorfe

## Abstract

KRAS is a high-value therapeutic target for the treatment of cancer. Two covalent inhibitors, sotorasib and adagrasib, which target a specific codon 12 mutation (G12C), have received accelerated approvals for clinical use. Studies of these inhibitors ushered in the development of new inhibitors such as MRTX1133, that had entered clinical trials as a KRAS (G12D)-selective, non-covalent inhibitor. However, the subsequent failure of sotorasib as monotherapy and the recent termination of an early-phase clinical trial for MRTX1133 indicates that developing clinically effective allele-specific KRAS inhibitors remains a challenge, and that there is a need for further evaluation of KRAS inhibition mechanisms. Here, we show that the reportedly KRAS (G12D)-selective MRTX1133 also binds to G12C mutant KRAS with high affinity and suppresses MAPK signaling in cancer cell lines harboring KRAS (G12C). Intriguingly, its effect on the proliferation of KRAS (G12C) cancer cells is context-dependent; MRTX1133 robustly inhibits the proliferation of the pancreatic cancer cell line MIA PaCa-2 as well as the tumor growth of MIA PaCa-2 mouse xenografts, but it has little effect in lung cancer cells. These findings, together with similar other recent reports, question if allele-specific KRAS inhibitors are truly selective and highlight the need for strategies that take into account tissue and context-specific processes.

**Significance Statement:** MRTX1133 is a reportedly selective, non-covalent inhibitor for the KRAS oncogene with a glycine-to-aspartate (G12D) mutation that is present in about 40% of pancreatic cancers. Despite the overwhelming preclinical success, the early-phase clinical trial of MRTX1133 was recently terminated with undisclosed results. Through our *in vitro* and *in vivo* studies, we discovered that MRTX1133 is also a potent non-covalent inhibitor of a glycine-to-cysteine (G12C) KRAS mutation that works in pancreatic cancer but not in lung cancer models. Our findings are consistent with other recent reports on the activity of MRTX1133 in non-G12D mutants and highlight challenges in developing true allele-specific KRAS inhibitors via non-covalent mechanisms while also accounting for tissue-specific effects.

## Introduction

Mutations in the *KRAS* gene drive tumorigenesis and immune evasion in many lethal cancers including pancreatic and lung cancers (1–3). Initial successes following decades of research saw two KRAS4B (hereafter KRAS) inhibitors entering the clinic in recent years to treat non-small cell lung cancer (NSCLC) patients with a glycine-to-cysteine mutation at codon 12 (G12C) of KRAS: sotorasib and adagrasib (2, 4–7). These inhibitors covalently bind to the nucleophilic thiol group of cysteine and ‘lock’ KRAS (G12C) in the GDP-bound off-state by preventing cycling to the GTP-bound on-state (5, 6). Although sotorasib monotherapy failed in a subsequent large-scale clinical trial for NSCLC (8), it was recently approved for combination therapy with panitumumab, an epidermal growth factor receptor (EGFR) inhibitor, for the treatment of metastatic colorectal cancer (CRC) (9). Moreover, adagrasib monotherapy showed better progression-free survival than chemotherapy in a recent clinical trial for KRAS (G12C) NSCLC patients (10). While KRAS (G12C) represents the majority of KRAS-driven lung cancers (39%), it is less common in other cancers (2, 11). For example, G12C represents only 3-4% of KRAS mutations in CRC and 1-2% in pancreatic ductal adenocarcinoma (PDAC) (12, 13). Instead, G12D is the most frequent KRAS mutation in PDAC (40%) and in CRC (21%) (13, 14). However, the development of covalent inhibitors of KRAS (G12D) is complicated by the fact that aspartate is a much weaker nucleophile than cysteine. The alternative was to develop G12D-selective non-covalent KRAS inhibitors. To this end, Wang et al. conducted an extensive modification of adagrasib to develop the G12D-selective, high-affinity, non-covalent KRAS inhibitor MRTX1133 (15).

MRTX1133 has since been a leading candidate in the race for KRAS G12D-selective inhibitors (15, 16). However, its Phase I clinical trial was recently terminated with undisclosed results (NCT05737706). Considering that an extensive panel of preclinical models consistently showed robust and durable responses to both MRTX1133 and sotorasib (4, 5, 15, 16), the recent clinical setbacks highlight the need for further evaluation of KRAS inhibition mechanisms and the determinants of therapeutic response. Nonetheless, the development and study of KRAS inhibitors with diverse modes of action has exponentially advanced our understanding of KRAS biology. Notable highlights include: (a) oncogenic KRAS can maintain nucleotide cycling and is not “locked” in the constitutively active state (17, 18); (b) loss of the GTPase function of oncogenic KRAS can be pharmacologically restored (19), (c) co-targeting wild-type mutant KRAS *in vivo* is feasible without systemic toxicity (20), and (d) maintenance of oncogenic signaling in PDAC is predominantly through the KRAS-RAF-ERK (MAPK) pathway (21). In the same spirit, our continuing investigation of KRAS inhibitors led to the unexpected finding that the ostensibly KRAS (G12D)-selective MRTX1133 also binds to and inhibits KRAS (G12C) in a PDAC cell and mouse cancer models. Combined with other recent reports (22, 23), this finding confirms that the selectivity of MRTX1133 arises not just from a salt-bridge with Asp12 (15) but also by eliciting an arrested GTPase cycling (24) and potentially from other mechanisms yet to be uncovered. These observations not only reveal new insights into KRAS inhibition but may also contribute to a potential explanation for the failure of MRTX1133 at its early-phase clinical trial.

## Results

### The KRAS (G12D)-selective inhibitor MRTX1133 inhibits growth of a G12C PDAC cell line

We have tested the inhibitory activity of MRTX1133 in 2D cell viability assays using six PDAC cell lines with distinct genetic backgrounds and harboring G12D, G12C, G12V or G12R KRAS mutations and one with KRAS wild type (WT) (**Figure 1A**). Consistent with previous reports (16, 22, 25), we found that MRTX1133 inhibits cell proliferation of the KRAS (G12D) mutant cell lines AsPc-1 and SW1990 (IC_50_ = 7-10 nM) but not the G12V PaTu 8902, G12R PSN-1, or the WT BxPc-3 PDAC cells (**Figure 1A, S1A-B**). Unexpectedly, however, we found that MRTX1133 inhibits the proliferation of the KRAS (G12C) PDAC cell line MIA PaCa-2 with an IC_50_ of 149 nM. To the best of our knowledge, neither the original report nor the subsequent pre-clinical testing of MRTX1133 included IC_50_ values for MIA PaCa-2 (15, 16). Other reports on the sensitivity of MIA PaCa-2 or other G12C cancer cell lines to MRTX1133 are conflicting (22, 23, 25, 26). Therefore, we repeated the experiments in Rasless mouse embryonic fibroblast (MEF) cells (**Figures 1B, S1C**), which lack endogenous H, N, and KRAS and therefore their growth depends only on an exogenously expressed KRAS or BRAF (27). In these cells, MRTX1133 is selective to KRAS (G12D) and does not inhibit the growth of other KRAS mutant or WT cell lines (**Figures 1B, S1C**). Notably, no significant inhibition was observed in the KRAS (G12C) MEF cells. The unexpected finding in the KRAS (G12C) PDAC cell line that is contradicted by data from an MEF cell line with the same mutation inspired us to further investigate the issue using binding experiments, additional KRAS (G12C) cancer cell lines, a 3D cell viability assay, and a MIA PaCa-2 xenograft mouse model.

**Figure 1.**
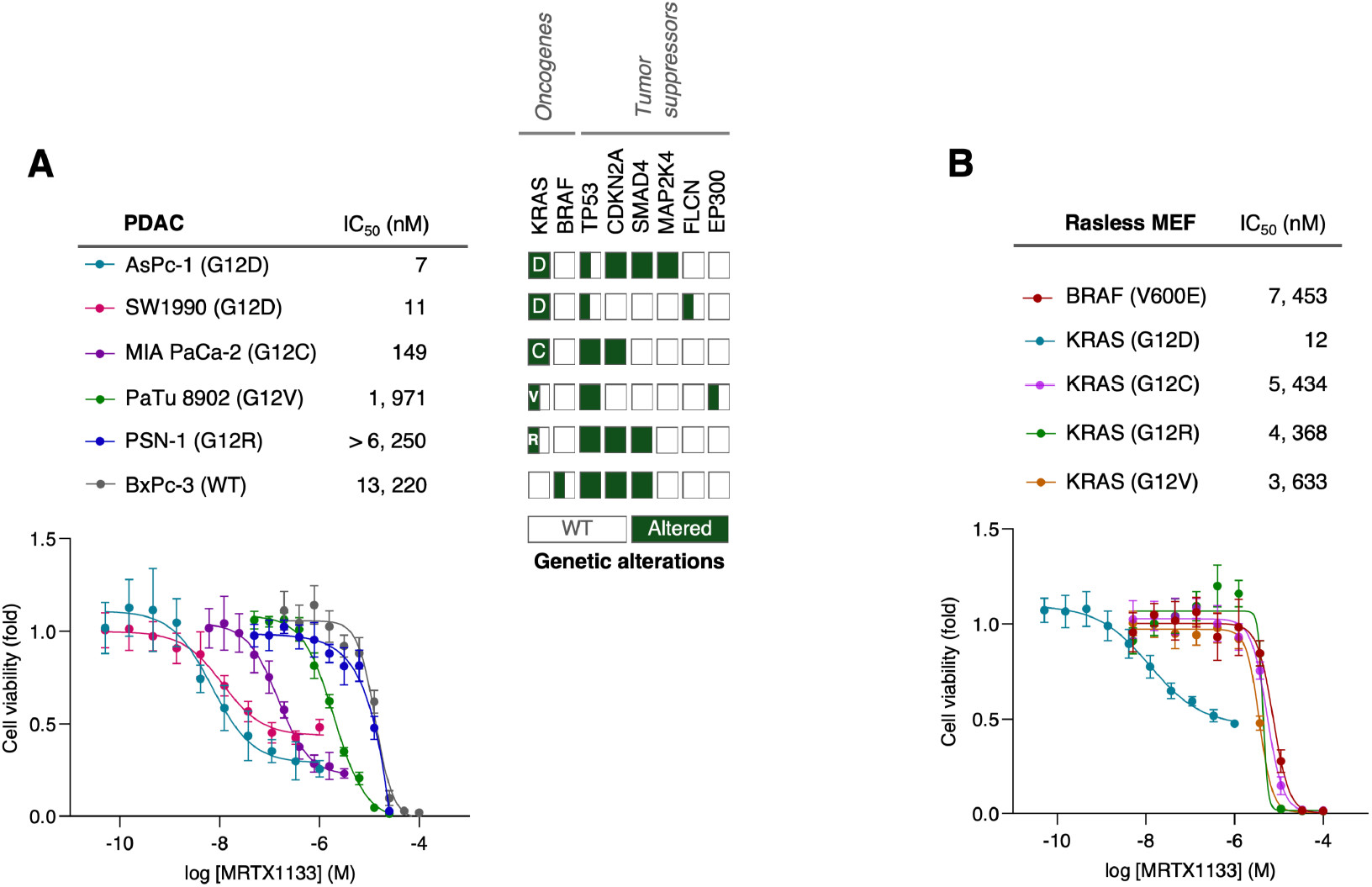
MRTX1133 inhibits cell proliferation of KRAS (G12D) and (G12C) PDAC but not MEF cell lines. The indicated human pancreatic ductal adenocarcinoma (PDAC) cell lines with KRAS (G12X) mutations (**A**) and isogenic and Rasless MEFs expressing human KRAS (G12X) or BRAF (V600E) mutations (**B**) were treated with MRTX1133 for 72 hours and cell viability was measured using CyQUANT^™^ assays. IC_50_ values were determined from curve fitting and are representative of two or three independent experiments (see Figures S1A-C for repeat experiments) with six different wells per concentration per experiment (n=6). Error bars represent standard deviation (SD). The tables in **A** indicate gene variants in each PDAC cell line, with full and half green-shaded boxes indicating homozygous and heterozygous alterations, respectively. Cell seeding densities: 2,000 cells/well for AsPc-1, SW1990, PaTu 8902, PSN-1; 4,000 cells/well for MIA PaCa-2, BxPc-3; 3, 000 cells/well for all Rasless MEFs.

### MRTX1133 directly binds to KRAS (G12C) with high affinity

While a key factor for the selectivity of MRTX1133 for KRAS (G12D) is likely a salt-bridge between its protonated bicyclic piperazinyl warhead and the Asp12 side chain (15), additional mechanisms of selectivity have been proposed (22, 24, 28). MRTX1133 was derived from the KRAS (G12C) covalent inhibitor MRTX849 (adagrasib) (**Figure 2A**), and molecular docking targeting the switch II pocket (originally called pocket p2) (29) of GDP-KRAS (G12C) yielded a highly favorable binding energy score of -13 kcal/mol and a similar binding pose to adagrasib (**Figure 2B**). The two inhibitors also engage a similar set of important KRAS residues, such as those described previously (22). Consistent with this prediction, we found using microscale thermophoresis (MST) that MRTX1133 binds to GDP-KRAS (G12C) with high affinity (K_D_ ∼ 0.3 nM) (**Figure 2C**), similar to nanomolar K_D_s found in other reports of MRTX1133 with GDP-KRAS (G12C) (30), as well as with G12V, G13D, Q61H/R, A146T, and WT (24, 30, 31). In the docked pose, the distance between the positively charged nitrogen on the piperazinyl moiety of MRTX1133 and the Cys12 thiol sulfur was 3.8 Å, suggesting the potential for a weak H-bonding interaction (**Figure 2B**).

**Figure 2.**
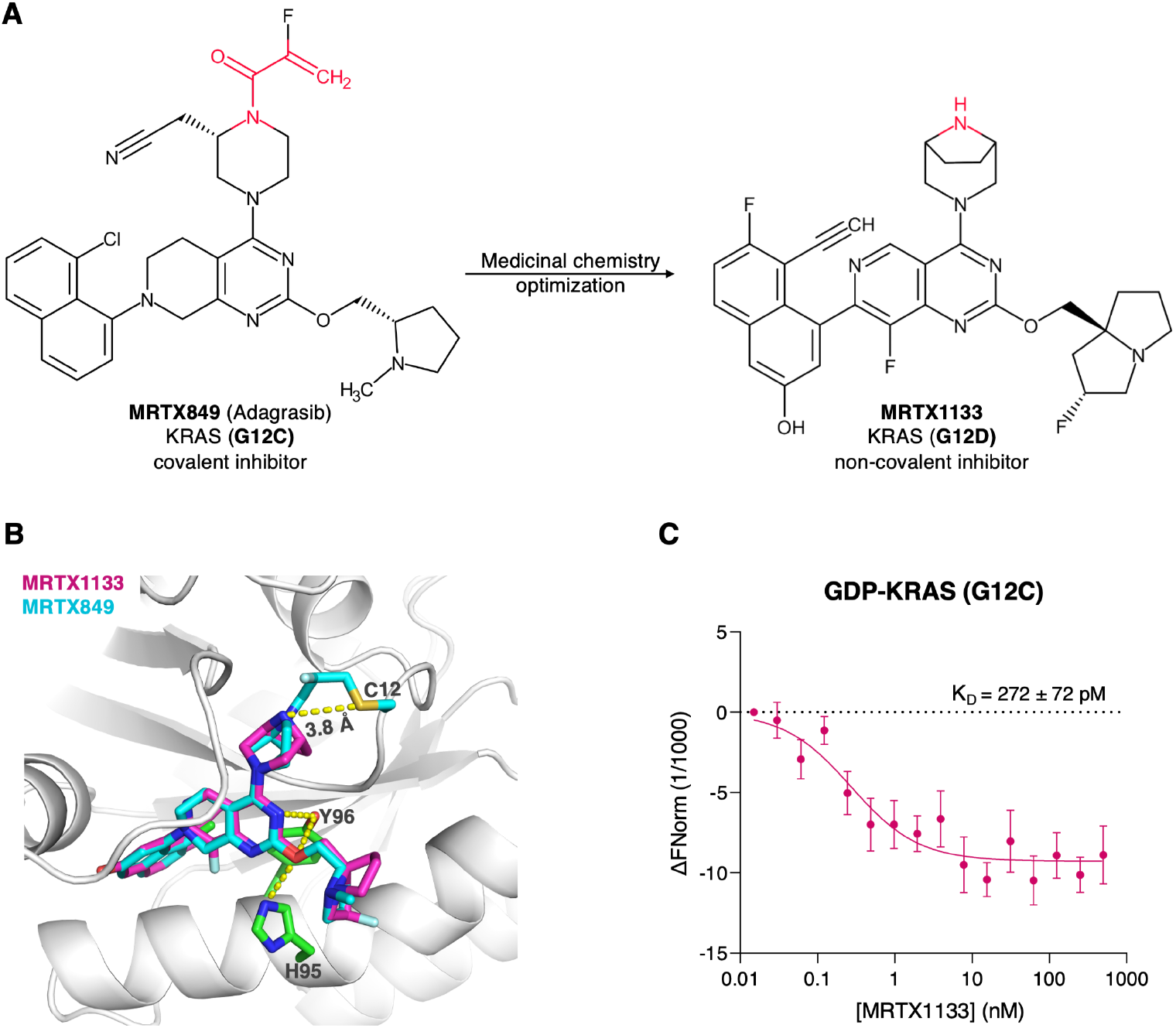
MRTX1133 binds to GDP-bound KRAS (G12C) with high affinity. (**A**) MRTX849, a KRAS (G12C) covalent inhibitor, was used as starting point for medicinal chemistry optimization to obtain MRTX1133, a non-covalent KRAS (G12D) inhibitor (15). Red highlights show acrylamide and bicyclic piperazinyl nitrogen warheads for G12C and G12D mutant specificity, respectively. (**B**) Molecular docking of MRTX1133 on GDP-bound KRAS (G12C) (PDB 6UT0) predicts strong binding affinity (Vina docking score -13 kcal/mol) and a similar binding pose to MRTX849 including shared H-bonding interactions (yellow dashed lines). (**C**) Microscale thermophoresis and calculated K_D_ for the interaction of recombinant GDP-KRAS (G12C) with MRTX1133 using three different readings from one capillary tube per concentration (n=3) (see Methods). Error bars = SD.

### MRTX1133 inhibits ERK phosphorylation but not proliferation of KRAS (G12C) lungcancer cells

As noted earlier, G12C is the dominant KRAS mutation in lung cancers (2, 11). Given the high affinity binding of MRTX1133 to recombinant KRAS (G12C) and its inhibition of a KRAS (G12C) PDAC cell proliferation, we wondered if it would also have an effect on KRAS (G12C) lung cancer cells. In a 2D cell viability assay format, we found that MRTX1133 has a small-to-negligible effect on the proliferation of the four lung cancer cell lines that we have tested, irrespective of zygosity or genetic background (**Figures 3A, S2A**). To check if this may be due to the 2D assay format, we repeated the experiments in 3D spheroids. We found that all four cell lines remained insensitive or barely sensitive to treatment by MRTX1133 (**Figure 3B**), consistent with the data from the 2D model. For comparison, we conducted the same 3D growth experiments in MIA PaCa-2 using adagrasib as a positive control. We found robust inhibition of MIA PaCa-2 proliferation after 3 days of treatment with an IC_50_ comparable to the 2D assay (**Figure 3C**), but with reduced effect with longer treatment time (**Figure S2B**) similar to previous findings in G12D models (16, 25). These results show that the inhibitory activity of MRTX1133 in the PDAC cell line MIA PaCa-2 but not in lung cancer cells with the same KRAS mutation is robust and not a consequence of experimental conditions.

**Figure 3.**
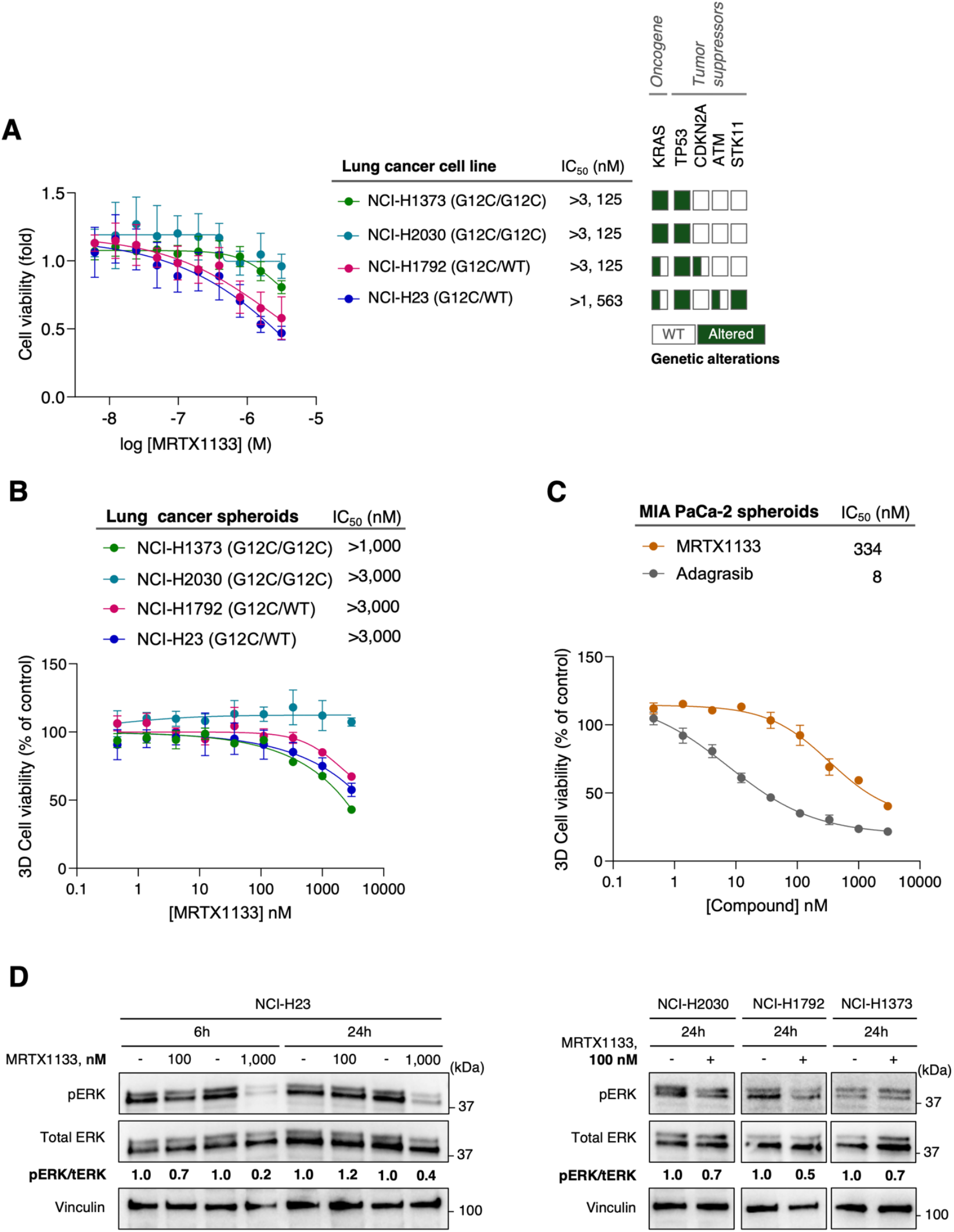
MRTX1133 has no effect on the proliferation of lung cancer cells but inhibits MAPK signaling. **(A)** Human lung cancer cell lines with KRAS (G12C) mutation were treated with MRTX1133 for 72 hours and cell viability was measured using CyQUANT^™^ assays. IC_50_ values were determined from curve fitting and are representative of two independent experiments (see Figure S2A for a repeat experiment) with six different wells per concentration per experiment (n=6). Cells were seeded at 2,000 cells/well. Error bars = SD. Shaded boxes in the respective tables indicate gene variants per cancer cell line (full – homozygous, half – heterozygous). **(B)** 3D spheroids of the same lung cancer cell lines treated with MRTX1133 for 3 days and cell viability was measured using CellTiter-Glo reagent. IC_50_ values were determined from curve fitting using three different wells per concentration seeded from one cell culture flask (n=3). Cells were seeded at 5, 000 cells/well. Error bars = SD. **(C)** The same as in **B** but for the MIA PaCa-2 cell line treated with MRTX1133 or adagrasib. IC_50_ curves are representative of two independent experiments (see Figure S2B for repeat experiments) with three different wells per concentration per experiment. **(D)** The lung cancer cell lines were treated with 100 nM or 1,000 nM of MRTX1133 for 6 hr or 24 hr and lysates were immunoblotted for pERK and total ERK with vinculin as loading control. The blots shown here are representatives of two independent experiments for each cell line.

Because oncogenic KRAS signaling primarily occurs through the MAPK pathway (21), we wondered if the difference in growth inhibition between the PDAC and lung cancer cells studied here may result from a differential effect on this pathway. We therefore conducted immunoblot analysis of ERK phosphorylation levels (pERK) to test for MAPK pathway inhibition. We found consistent and significant reduction of pERK levels upon treatment of the lung cancer cells for 6 hrs or 24 hrs with either 100 nM or 1,000 nM of MRTX1133 (**Figure 3D**). Similarly, treatment of the PDAC MIA PaCa-2 for 3, 6 and 24 hrs with MRTX1133 shows a robust, dose-dependent, and durable inhibition of this pathway, with significant decreases in pERK levels even at 100 nM (**Figures 4A, S3A**). Remarkably, very similar inhibitory profiles were observed when comparing pERK levels in MIA PaCa-2 cells treated with 100 nM MRTX1133 or the covalent KRAS (G12C) inhibitors sotorasib and adagrasib (**Figures 4B, S3B**). In fact, there are no significant differences in pERK levels among the three inhibitors at 3 hrs of treatment, and only small differences at other incubation times (**Figure 4C**). Furthermore, the inhibitory activity of both MRTX1133 and the covalent inhibitors is reduced upon extended incubation (50% inhibition at 24 hrs), suggesting a shared mode of inhibition. These data show that MRTX1133 not only effectively binds to KRAS (G12C) but also suppresses MAPK signaling in the two tissues that we have tested. Taken together, the reduction of pERK levels in all cell lines that we have studied is consistent with the high-affinity binding of MRTX1133 to KRAS (G12C) and the growth inhibition in the KRAS (G12C) PDAC cell line, while the lack of growth inhibition in the KRAS (G12C) lung cancer cells likely involves regulatory processes beyond the KRAS GTPase cycle.

**Figure 4.**
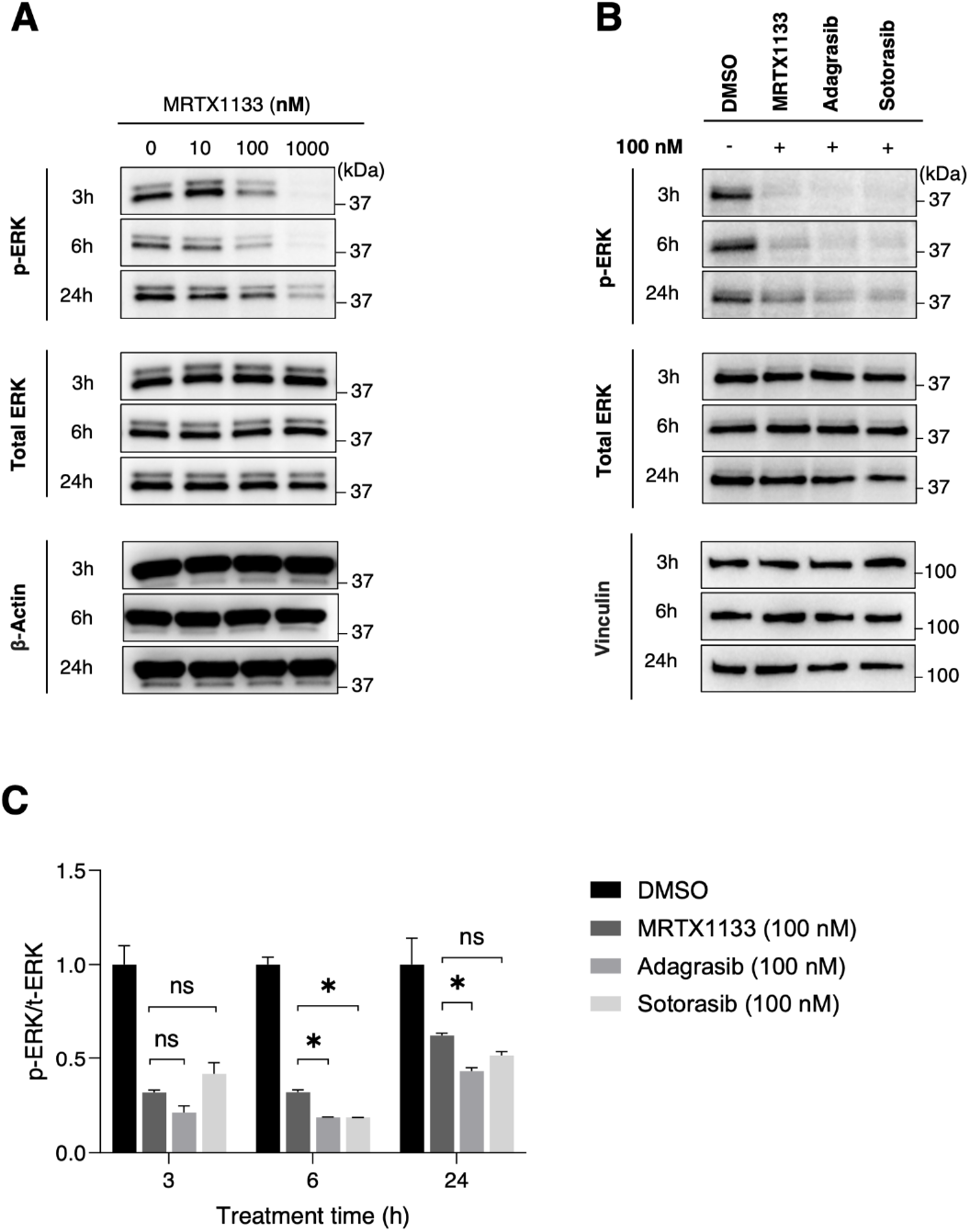
The non-covalent inhibitor MRTX1133 is as potent as the covalent KRAS (G12C) inhibitors sotorasib and adagrasib in reducing MAPK signaling in MIA PaCa-2 cells. MIA PaCa-2 cells were treated with indicated concentrations of MRTX1133 **(A)** or 100 nM of MRTX1133, adagrasib and sotorasib (**B** and **C**) for 3, 6, 24 hours and cell lysates were harvested and immunoblotted for pERK and total ERK. Shown in panels **A** and **B** are representative western blots from two independent experiments (see Figures S3A, B for repeat experiments), and changes in pERK levels shown in **C** were quantified relative to total ERK levels and normalized to untreated (DMSO) samples. *p<0.05 on ordinary one-way ANOVA. n.s. = not significant. MRTX1133, adagrasib, and sotorasib have significantly lower values (p<0.05) than DMSO control in all time points (asterisks not shown).

### MRTX1133 inhibits tumor growth in a MIA PaCa-2 xenograft mouse model

To test if our observations in binding, cell signaling, and 2D as well as 3D cell proliferation assays translate into anti-tumor efficacy in mice, we tested the ability of MRTX1133 to inhibit MIA PaCa2 xenograft tumor growth. **Figure 5A** shows that MRTX1133-treated mice exhibited a significantly lower rate of tumor growth compared to vehicle control (captisol). Importantly, there was no significant difference in weight of the mice throughout the treatment, indicating a lack of toxicity (**Figure 5B**). Consistent with effects in tumor growth, MRTX133-treated mice had significantly reduced tumor burden at the end of the experiment (**Figures 5C, D**). This is a remarkable effect for an inhibitor originally thought to be ineffective for KRAS (G12C) (25, 26). Moreover, in the current experiment, the inhibitor was administrated at 30 mg/kg dose, which is likely insufficient for maximal effect. While this dose was chosen based on literature reports on KRAS (G12D) mouse models (16, 25, 26), it did not account for the ∼15-fold difference in IC_50_ between G12D and G12C (**Figure 1A**). Also, MRTX1133 may work even better in immunocompetent mice since its activity is augmented by T-cells in the tumor microenvironment (25).

**Figure 5.**
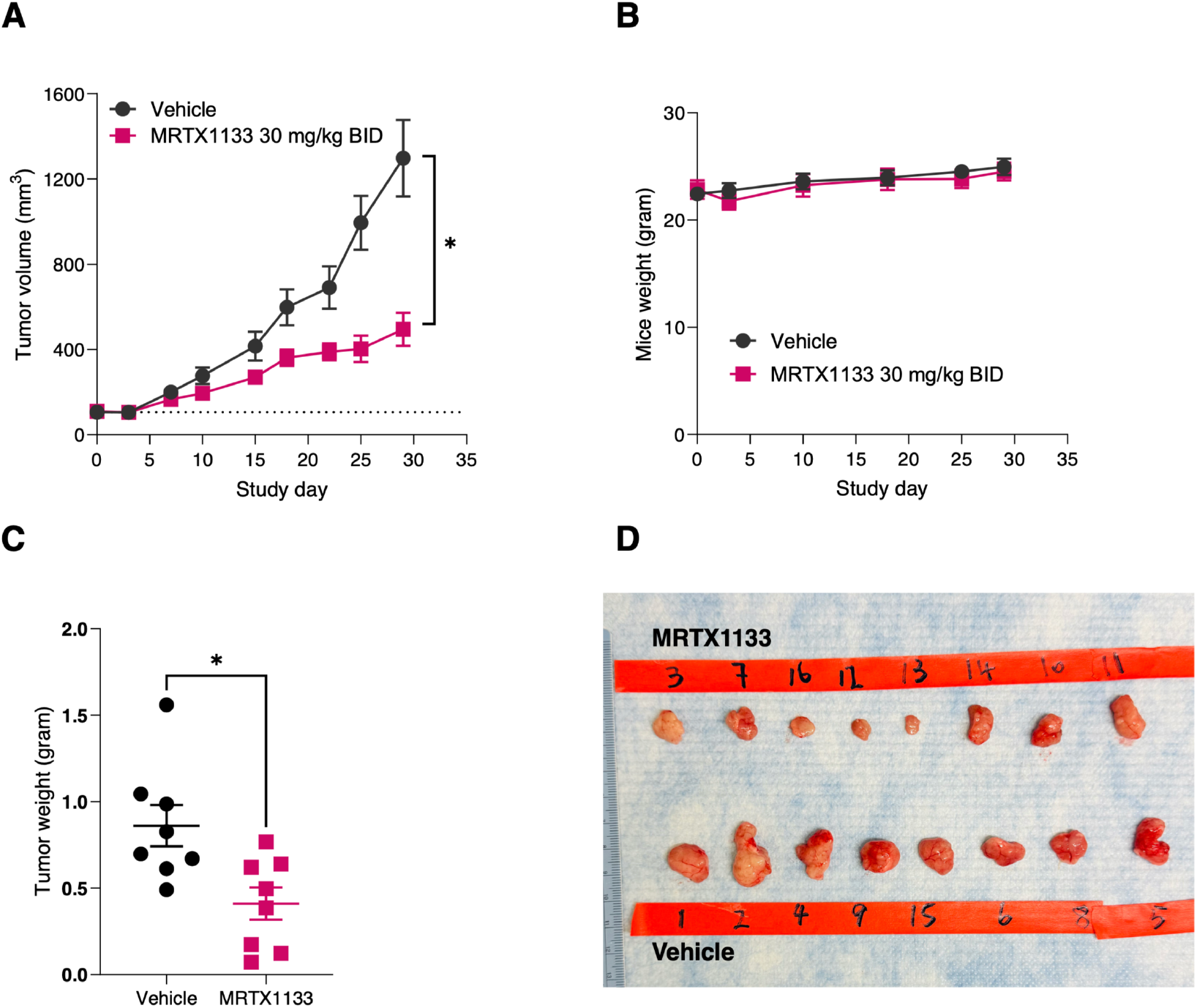
MRTX1133 inhibits MIA PaCa-2 in mouse xenografts. (**A**) Mice bearing MIA PaCa-2 xenografts were treated twice a day (BID) by intraperitoneal administration of 30 mg/kg MRTX1133 (n=8) or vehicle (n=8) for 28 days. (**B**) Mice were weighed periodically during the treatment duration. (**C**) Tumors were harvested and weighed on the 29^th^ day. (**D**) Harvested tumors. Error bars in **A-C** represent standard error of the mean (SEM); *p<0.05 on two-tailed, unpaired t-test.

## Discussion

KRAS is a critical oncogene driver in many lethal human cancers (2, 7, 16, 20). Recent discoveries of covalent and non-covalent KRAS inhibitors for the treatment of cancer have ignited excitement in the RAS community and beyond (32). However, recent reports of setbacks regarding two of the most highly promising inhibitors, sotorasib and MRTX1133, as KRAS monotherapies raised new questions about our understanding of KRAS biology and its inhibition mechanisms (2, 4, 8, 16). Among these questions are whether allele-specific inhibitors are truly selective and how significant the potential impact of tissue and context-dependent inhibition might be. Our finding that MRTX1133, which was thought to be a highly specific KRAS (G12D) noncovalent inhibitor, is also a potent inhibitor of some but not all KRAS (G12C)-driven cancers reinforces the significance and timeliness of these questions.

The high-affinity binding of MRTX1133 to KRAS (G12C) (**Figure 2**) may be attributed to its overall structural similarity to its parent G12C covalent inhibitor adagrasib and its ability to engage the same set of KRAS residues (e.g., His95, Tyr96) (22, 33). As in the covalent G12C-specific inhibitors adagrasib and sotorasib (5, 6), MRTX1133 preferentially binds to GDP-bound than GTP-bound mutants and WT KRAS (24, 31). The primary mode of inhibitory action of the covalent inhibitors has been shown to involve impairing GDP/GTP exchange (4, 7), and we found that MRTX1133 inhibits GDP/GTP exchange in KRAS (G12C) to the same extent as adagrasib and sotorasib (**Figure S4**). This suggests that this mode of action contributes to the ability of MRTX1133 to suppress MAPK signaling in the KRAS (G12C) mutant cell lines that we have tested (**Figures 3D, 4**), and its effect in MIA PaCa-2 proliferation in both monolayer and spheroid models as well as in athymic nude mouse xenografts (**Figures 1A, 3A-C, 5**). Although the lack of other established human KRAS (G12C) PDAC cells prevents generalization, our findings are supported by previous observations on the activity of MRTX1133 in other non-G12D KRAS mutants (16, 22– 24, 30, 31), and support a previous suggestion that Asp12 is important but not required for the activity of MRTX1133 (22, 28). Together, these results show that true allele-specificity remains a challenge for directly inhibiting KRAS through non-covalent mechanisms.

The apparent “pan-KRAS” activity of MRTX1133 coupled with its enhanced activity towards KRAS (G12D) would have been grounds for optimism regarding its success in the clinic. While the primary reason for the termination of the clinical trial is not fully fully clear, the heterogeneity of cellular responses to MRTX1133 in KRAS (G12C) PDAC versus lung cancer and MEF cells observed in this work suggests roles for tissue-specific factors. Other evidence supporting this notion include the following. In KRAS (G12D) cell line-or patient-derived tumor xenograft studies of MRTX1133, tumor regression was seen in 8/11 PDAC, 2/8 CRC, and 0/4 lung cancer models (16), suggesting a context-dependent inhibitory activity. Similar to our findings in KRAS (G12C) lung cancer cell lines, a study in KRAS (G12D) patient-derived PDAC cell lines showed that MRTX1133 inhibits ERK phosphorylation but not cell proliferation, and co-targeting of the mammalian target of rapamycin (mTOR) overcame this apparent resistance (26). Depending on KRAS mutation and tissue origin, tumors can also show variations in treatment-associated differentially expressed genes, ERK-independent bypass and feedback pathways, or response to combination therapies involving other KRAS pathway inhibitors (7, 16). Taking these findings into account, we propose that the differential effect on cancer cell growth we observed suggests possible tissue-specific bypass of the MAPK pathway in the lung but not PDAC cancer cells or different feedback mechanisms operating in the two tissues (7, 16). Further, a recent study showed that MRTX1133 could be a “double-edged sword” in that it stabilizes KRAS in the GDP state by impeding activation and in the GTP state by reducing both intrinsic and GAP-mediated hydrolysis (24). BRAF-binding to the MRTX1133-bound GTP-KRAS (G12D) restored the intrinsic hydrolysis, albeit incompletely (24). We surmise that other key signaling players (e.g., RAF, mTOR) may be mediating the heterogeneity of responses to MRTX1133 in different KRAS-mutant cancers with unique genetic and immunologic makeup. Taken together, irrespective of its fate in clinical trials, MRTX1133 continues to advance our understanding of KRAS biology and to provide key lessons that will inform future efforts toward broad-acting KRAS inhibitors.

## Conclusion

The recent setbacks to initial breakthroughs in allele-specific KRAS inhibition underscores ongoing challenges in developing effective clinical therapies for KRAS-driven cancers. In this work, we have shown that the KRAS (G12D)-selective inhibitor MRTX1133 is also a non-covalent inhibitor of KRAS (G12C). MRTX1133 strongly binds to GDP-KRAS (G12C) and inhibits MAPK signaling in KRAS (G12C) cancer cell lines as assessed by levels of ERK phosphorylation. As in covalent G12C inhibitors, MRTX1133 impairs SOS1-mediated nucleotide exchange and thus KRAS activation. These findings are consistent with previous reports of the activity of MRTX1133 in non-G12D mutants that lack the Asp12 selectivity handle, including G12C. However, MRTX1133 inhibits tumor growth and proliferation of a KRAS (G12C) PDAC cell line but has no effect in lung cancer cells harboring the same KRAS mutation. This lack of growth inhibition in lung cancer cells despite reduced ERK phosphorylation and the robust effect in the PDAC cell line suggest possible ERK-independent bypass mechanisms or context-dependent effects and highlight the challenge of allele-specific KRAS inhibition via non-covalent mechanisms.

## Materials and Methods

### Cell culture and reagents

Human pancreatic ductal adenocarcinoma (PDAC) cell lines AsPc-1, SW1990, and PaTu-8902 were grown in DMEM (Gibco 12100-046) supplemented with 10% FBS (Corning 35-011-CV) and 100 units/mL of penicillin and 100 µg/mL of streptomycin (Cytiva SV30010). BxPc-3, PSN-1, and lung cancer cell lines NCI-H23, NCI-H1292, H1373, H2030 were grown in RPMI 1640 medium (Gibco 31800-022) supplemented with 10% FBS and 100 units/mL of penicillin and 100 µg/mL of streptomycin. MIA PaCa-2 cells were grown in DMEM + GlutaMAX^™^ (Gibco 10569-010) supplemented with 10% FBS. Rasless mouse embryonic fibroblasts (MEF) cells expressing different KRAS mutants were grown in DMEM supplemented with 10% FBS and 4 µg/mL of blasticidin (ChemCruz F2422) to maintain the stable expression of mutant KRAS. In all cases, cells were maintained in a humidified 5% CO_2_ incubator at 37° C. MRTX1133 (Chemietek Lot #3), MRTX849 (adagrasib) (Chemietek Lot#5) and AMG-510 (sotorasib) (MedChemExpress HY-114277) were prepared at 10 mM stock concentration in DMSO.

### Cell proliferation assays

2D cell proliferation assays were conducted using CyQUANT^™^ cell proliferation assay kit (Invitrogen^™^) following manufacturer’s instructions. Cells were plated in 96-well, black-walled, clear-bottom plates (Corning 3903) at 2,000 – 4,000 cells/well. After 24 hours of seeding, cells were treated with serial dilutions of MRTX1133 (with a final DMSO concentration of 0.5%) for 72 hours. Growth media were then removed, and adherent cells were washed with PBS before freezing for at least 24 hours at -80° C. Plates were thawed and CyQUANT dye (Invitrogen C7026 A) solution was prepared in the kit’s lysis buffer (Invitrogen C7026 B), and then 200 µL of solution was added to each well for a five-minute incubation. Fluorescence was measured in Tecan Infinite M200 plate reader (excitation: 488 nm; emission: 525 nm). For 3D cell proliferation assay (7, 16, 20), the CellTiter-Glo (CTG) reagent was used following manufacturer’s protocol. MIA PaCa-2 cells were seeded at 5,000 cells/well in ultra-low attachment plates (Thermo Fisher Scientific CN163320) and centrifuged at 1,000 rpm for 10 min. After 24 hours, cells were treated with serial dilutions of MRTX1133 for 72 hours. Cell viability was determined using CTG reagent and chemiluminescence was measured using Tecan Infinite M200 plate reader. For both 2D and 3D, cell viability was determined by normalizing fluorescence or chemiluminescence units per concentration to DMSO control and IC_50_ values were calculated through curve fitting with dose-response-inhibition – variable slope (four parameter) on GraphPad Prism 10.0. Viable cells were counted in a cytometer (Invitrogen 100078809) using Trypan blue (Invitrogen T10282) to seed the same number of viable cells per well at the beginning of the assay.

### Binding assay and molecular docking

Protein expression and purification and determination of protein-ligand dissociation constant (K_D_) using microscale thermophoresis (MST) were as described in previous report (34-36). Briefly, recombinant GDP-bound KRAS (G12C) was fluorescently labeled following the manufacturer’s protocol using Alexa Fluor 647 Dye (Thermo Fisher Scientific). Labeled KRAS was purified using the kit column (NanoTemper) and diluted to 100 nM with MST assay buffer (40 mM HEPES, pH 7.5, 5 mM MgCl_2_, 100 mM NaCl + 0.05% Tween-20). Then, serial dilutions of MRTX1133 were made in MST assay buffer + 2-4% DMSO and added to equal volumes of the protein. The mixtures were incubated for at least 30 minutes and loaded in capillaries for measurement in Monolith NT.115 using 20% LED power and 40% MST power. To determine K_D_, curve fitting was done using one-site specific binding model in GraphPad Prism 10.0. For site-directed molecular docking, the virtual screening software package Vina (37). implemented in PyRx (https://pyrx.sourceforge.io/) was used to dock MRTX1133 (prepared from PDB 7RPZ) to the KRAS (G12C) structure PDB 6UT0 with search space defined by residues V8, G10, D12, T58, G60, E62, Y64, R68, D69, and H95 (15). Protein-ligand interactions were examined using PyMOL and Protein Ligand Interaction Profiler (PLIP) (38).

### Cell signaling assays

MIA PaCa-2 cells were plated in 6-cm dish (GenClone 25-202) until about 40-60% confluency and treated with fresh media containing MRTX1133 or DMSO (Fisher Bioreagents BP231-100) for a specified period (3, 6, or 24 hrs). Then, the growth medium was discarded, and cells were collected with cold PBS (Fisher BioReagents BP399-1), pelleted by centrifugation at 21,000 *x g*, 40 min, 4° C, and lysed in 400 µL lysis buffer (Thermo Scientific, #1862301) containing protease and phosphatase inhibitor cocktail (Roche), pH 7.4. Total protein concentration in cell lysates was determined by BCA assay (Thermo Scientific 23228, A, 1859078, B). Cell lysate was mixed with sample buffer (GenScript MB01015) and denatured at 95° C for 5 min. Equal amounts (20 µg) of protein were separated by SDS page in a 10% SDS-PAGE gel (Bio-Rad) and then analyzed by Western blotting using PVDF membrane. Membranes were blocked in 5% blocking milk (RPI M17200-500.0) in TBS (BioRad #1706435) with 0.1% Tween-20 (Sigma P1379-1L) for 1 h and then overnight with primary antibody at 4° C. Membranes were then washed for 5 mins in TBS containing 0.1% Tween-20 (Sigma, #P1379-1L) three times, then hybridized with horseradish peroxidase (HRP)-conjugated secondary antibody in TBST for 1 h at room temperature. TBST washing was done for 5 min three times. Primary antibody dilution was 1:1000 for p-ERK (Cell Signaling Technologies #4370S), t-ERK (CST 910S), β-actin (Sigma A1978), and vinculin (CST, #13901S) while 1:3000 for secondary antibody (Goat anti-rabbit, CST #7074S or Goat anti-mouse, Invitrogen #G21040). Blots were developed by enhanced chemiluminescence (Pierce, #32106), imaged with Azure biosystems 280 imager, and quantified using Image J software. After pERK blots, membranes were stripped at room temperature for 30 min with gentle shaking in 1X stripping buffer (Thermo Scientific #W28K536) to reprobe for tERK and then stripped once more for β-actin.

### *In vivo* studies

3.0 x 10^6^ MIA PaCa-2 cells were resuspended in 50% Matrigel/50% PBS and injected subcutaneously into the hind flank of 6-week-old female outbred athymic nude mice (J:Nu cat #007850). Tumor volumes were measured with electronic calipers and volumes were calculated using the formula: volume = (length x width^2^)/2. When tumor volumes reached an average of ∼100 mm^3^, mice were randomized into two groups of eight (control and treatment). Mice were treated twice a day, 5 days a week for 4 weeks with MRTX1133 diluted in vehicle [10% Captisol (Cydex NC-05E-21045) in 50 mM citrate buffer, pH 5.0 (Teknova Q2443)] at a dose of 30 mg/kg via intraperitoneal injection. The control group received just the vehicle. The 30 mg/kg dose was chosen based on previous reports using KRAS (G12D)-mutant mouse models (16, 25, 26). The mice were weighed once a week and tumor volumes were measured twice a week. Mice were euthanized when tumor volumes reached 1,500 mm^3^ and the tumors were excised for further analysis.

## Supporting information

https://drive.google.com/file/d/1dPhAl0rlgudcTOuJgpahS0UOLevYoMVc/view?usp=sharing

## Data, Materials, and Software Availability

All study data are included in the article and/or SI Appendix.

## Acknowledgment

This work was supported in part by grants from the Cancer Prevention and Research Institute of Texas (CPRIT) grant # RP240104 and the National Institute of Health Institute of General Medicine (NIGMS) grant # R01GM144836. ACS acknowledges financial support from the Department of Science and Technology-Philippine Council for Health Research and Development.

