## Supplementary material for "MRTX1133 is a potent non-covalent KRAS (G12C) inhibitor with tissue-specific activity": https://drive.google.com/file/d/1dPhAl0rlgudcTOuJgpahS0UOLevYoMVc/view?usp=sharing

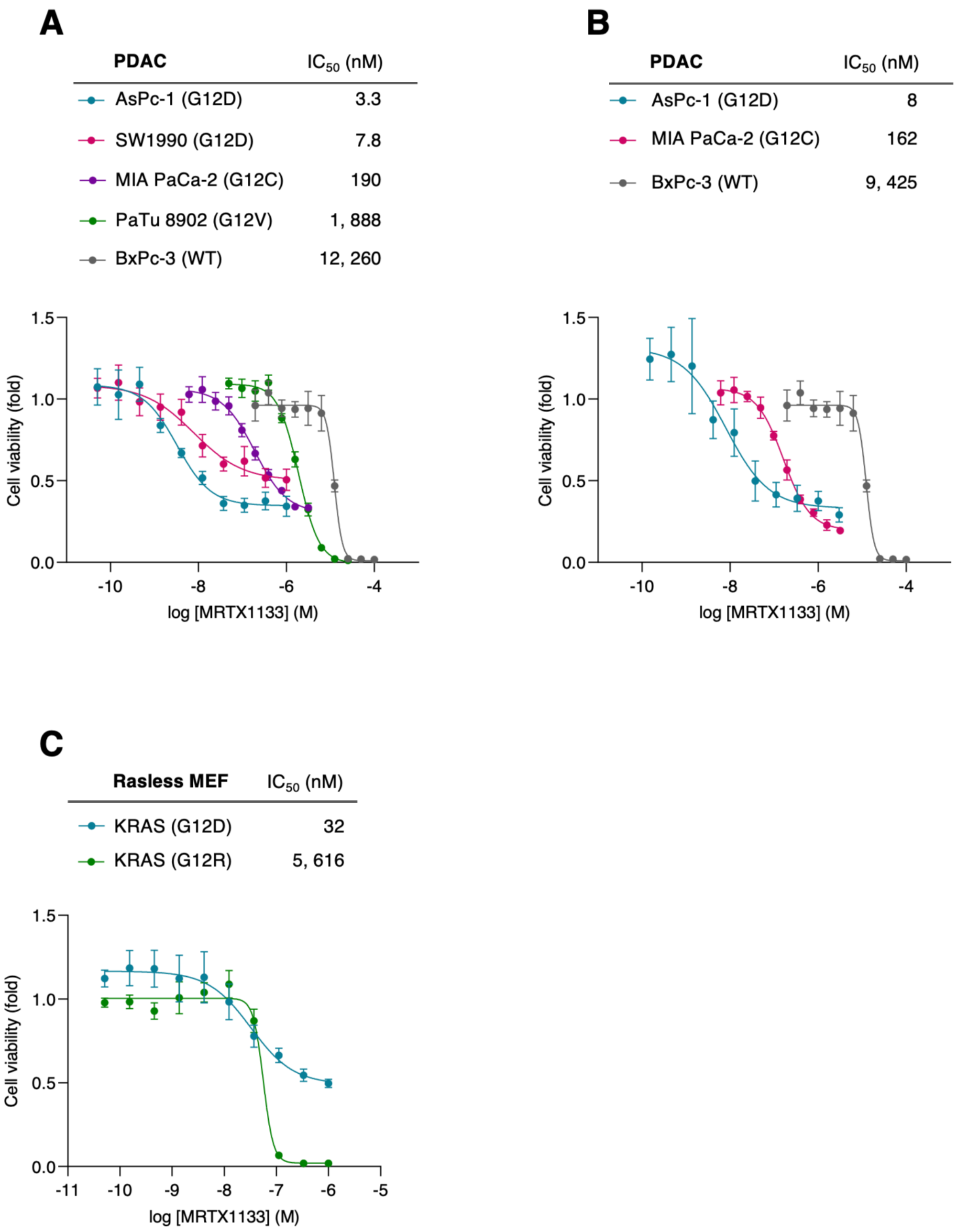


**Figure S1. MRTX1133 inhibits cell proliferation of KRAS (G12D) and (G12C) PDAC but not MEF cell lines.** The indicated human pancreatic ductal adenocarcinoma (PDAC) cell lines with KRAS (G12X) mutations (**A, B**) and isogenic and Rasless MEFs expressing human KRAS (G12X) or BRAF (V600E) mutations (**C**) were treated with MRTX1133 for 72 hours and cell viability was measured using CyQUANT$^{TM}$ assays. $IC_{50}$ values were determined from curve fitting using six different wells per concentration per experiment (n=6). Error bars = SD. Cell seeding densities in **A**: 2,000 cells/well for AsPc-1, SW1990, PaTu 8902; 4,000 cells/well for MIA PaCa-2, BxPc-3; in **B**: 2, 000 cells/well for all PDAC cell lines; in **C**: 3, 000 cells/well for all Rasless MEFs.

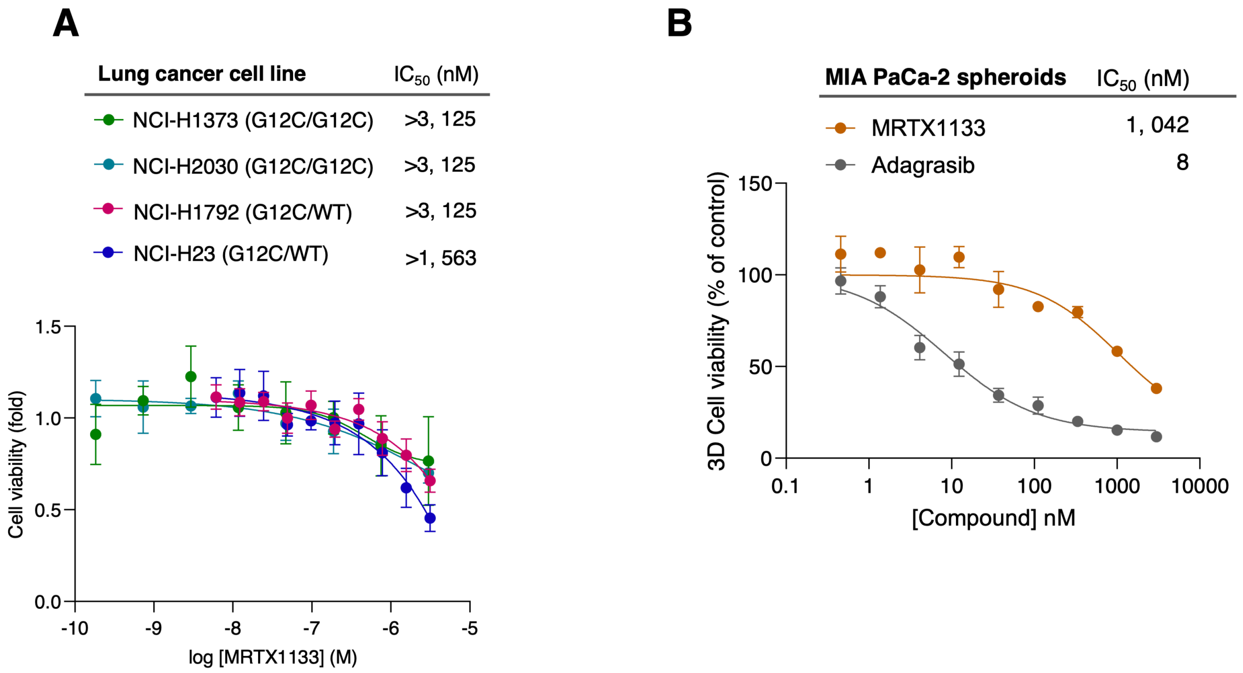


**Figure S2. MRTX1133 has no effect on the proliferation of lung cancer cells. (A)** Human lung cancer cell lines with KRAS (G12C) mutation were treated with MRTX1133 for 72 hours and cell viability was measured using CyQUANT™ assays. $IC_{50}$ values were determined from curve fitting using four-six different wells per concentration per experiment. Cells were seeded at 2,000 cells/well for NCI-H1373 and NCI-H2030 while 4,000 cells/well for NCI-H1792 and NCI-H23. Error bars = SD. **(B)** 3D spheroids of MIA PaCa-2 cell line treated with MRTX1133 or adagrasib for 8 days and cell viability was measured using CellTiter-Glo reagent. $IC_{50}$ values were determined from curve fitting (constraint, top=100) using three different wells per concentration per experiment (n=3). Cells were seeded at 5, 000 cells/well. Error bars = SD.

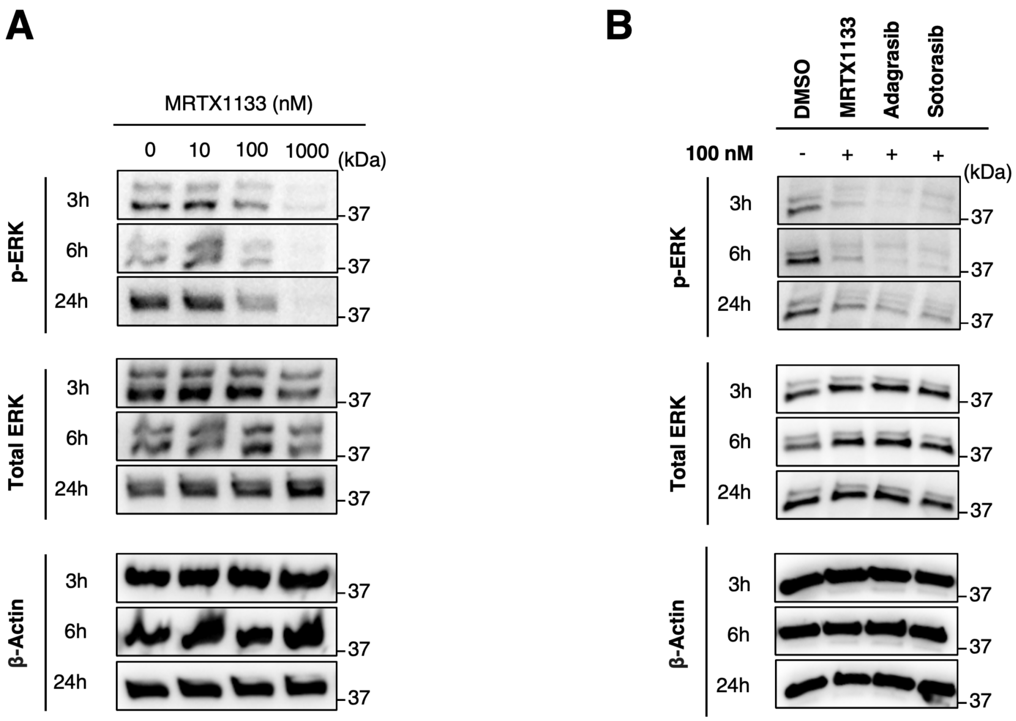


**Figure S3. The non-covalent inhibitor MRTX1133 is as potent as the covalent KRAS (G12C) inhibitors sotorasib and adagrasib in reducing MAPK signaling in MIA PaCa-2 cells.** MIA PaCa-2 cells were treated with indicated concentrations of MRTX1133 **(A)** or 100 nM of MRTX1133, adagrasib and sotorasib (**B**) for 3, 6, 24 hours and cell lysates were harvested and immunoblotted for pERK and total ERK.

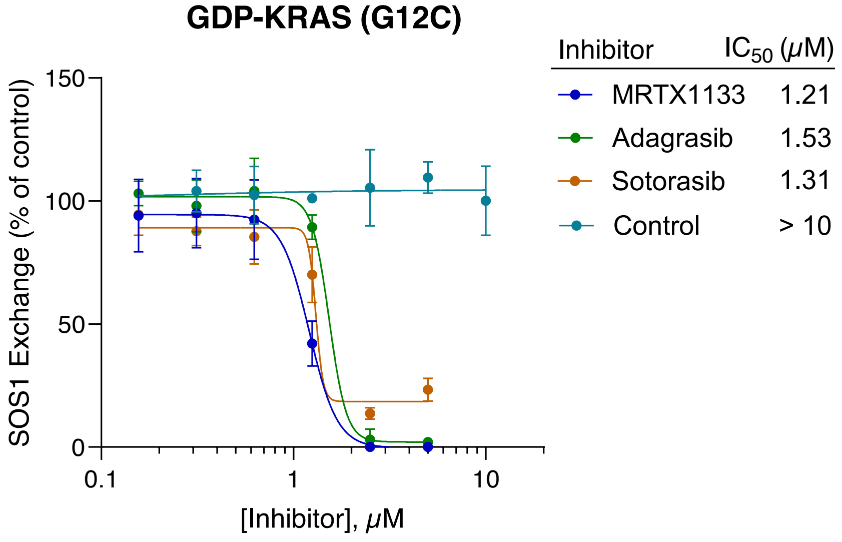


**Figure S4. MRTX1133 inhibits GDP/GTP exchange in KRAS G12C.** As described in our previous work (1, 2), rates of nucleotide exchange mediated by the RAS GEF SOS1 catalytic domain (residues 564-1049) were determined by monitoring the fluorescence intensity increase of BODIPY-GTP FL (Invitrogen #3057037) as it is exchanged with GDP in KRAS for 30 min in a buffer containing 25 mM Tris (pH 7.5), 50 mM NaCl, 10 mM MgCl2, and 1 mM TCEP. Varying concentrations of inhibitors were added with the reaction mixture (100 µL) containing 0.5 µM each of GDP-KRAS (G12C), BODIPY-GTP and SOS1; BODIPY-GTP was added just before measurements. Fluorescence intensities were read at 520 nm (excitation at 485 nm) using Tecan Infinite M200 microplate reader. Rates at every concentration were determined using dissociation-one phase exponential decay fitting in GraphPad Prism 10.0 and were normalized to DMSO. The data shows nucleotide exchange of recombinant GDP-KRAS (G12C) monitored by fluorescence increase of BGTP as it is exchanged with GDP by the catalytic domain of SOS1. Measurements were made in 3 different wells per concentration of MRTX1133, and with the G12C covalent inhibitors adagrasib and sotorasib as positive controls and an unpublished compound as a negative control. Rates at every concentration were normalized to DMSO and IC$_{50}$ was obtained as described previously (1, 2). Error bars = SD.
